# Boundary Homogenization and Numerical Modeling of Solute Transport Across the Blood–Brain Barrier

**DOI:** 10.1101/2025.07.01.662658

**Authors:** Reza Yousofvand, Gregory Handy, Jeffrey Tithof

**Author notes:** G.H. and J.T. contributed equally to this work. Correspondence (G.H.), (J.T.).

## Abstract

Effective clearance of amyloid-β (Aβ) from the brain is essential for preventing neurodegenerative diseases such as Alzheimer’s. A significant portion of this clearance occurs through the blood–brain barrier (BBB) via receptor-mediated transport. However, current models fail to capture the complex kinetics and spatial heterogeneity of receptors at the BBB. In this study, we derive a novel boundary condition that accounts for finite receptor kinetics, receptor density, and bidirectional transport across the BBB. Specifically, we develop a nonlinear homogenized boundary condition that ensures mass conservation and incorporates receptor-mediated Michaelis–Menten kinetics. We then implement this boundary condition in a cylindrical geometry representing a capillary surrounded by brain tissue. After verifying that the model matches an analytical steady state solution that we derive and that it yields realistic blood Aβ concentrations, we explore how realistic variations in parameter values drive changes in both steady state Aβ concentration and transient dynamics. Simulations and analytical results reveal that Aβ concentrations in the brain are sensitive to receptor number ratios, while concentrations in the blood are primarily affected by the blood clearance rate. Additionally, we use the model to investigate Aβ clearance during sequential sleep cycles and due to a pathological phenomenon, spreading depolarization. This work presents the first biophysically consistent boundary condition for Aβ transport across the BBB, offering a powerful tool for studying brain waste clearance under both physiological and pathological conditions.

## I. INTRODUCTION

An inevitable outcome of cellular metabolism is the generation of waste products. In most tissues, interstitial fluid containing these cellular byproducts drains into lymphatic vessels, which transport the fluid to the venous bloodstream. From there, the waste is distributed to various organs and eventually cleared from the body. However, the lymphatic system is absent in the brain. Being a highly active organ, the brain produces waste proteins such as amyloid-β (Aβ) that require clearance. If not cleared effectively, Aβ accumulates in the brain and results in development of diseases such as Alzheimer’s and related dementias [1–3]. Other than metabolic degradation, Aβ is cleared through the blood-brain barrier (BBB) to be carried away via blood flow or through the glymphatic system. The glymphatic system is a brain-wide clearance pathway in which cerebrospinal fluid (CSF) flows through perivascular spaces (PVSs) surrounding penetrating vessels, facilitating waste removal from the brain via regulated transport across astrocyte endfeet [4–9].

In humans, 50% of all generated Aβ in the brain is eliminated by proteolytic degradation and from the remaining 50%, approximately half is cleared through CSF flow, and the other half is cleared through the BBB [5]. Similarly, in mice, about 27% of total Aβ clearance is mediated by the BBB [10]. A higher contribution of BBB in clearance has been reported using the brain efflux index method, which indicated that 62% of intracerebrally-injected Aβ appeared in the blood [11]. These measurements illustrate the significance of the BBB in clearing Aβ from the brain and thus highlight its role in preventing the development of neurodegenerative diseases related to Aβ accumulation.

The BBB is a highly selective semipermeable border of endothelial cells around blood vessels that tightly regulates the movement of ions, molecules, and cells between the blood and the brain. For Aβ, two so-called “active transporters” exist, the receptor for advanced glycation end products (RAGE) and low-density lipoprotein receptor related protein-1 (LRP1) [12, 13]. LRP1, expressed on the abluminal side of brain endothelial cells, facilitates the efflux of Aβ from the brain into the cardiovascular circulation. This receptor-mediated transport is essential for reducing Aβ accumulation in the brain parenchyma [14–16]. Conversely, RAGE, located on the luminal side of endothelial cells, mediates the influx of circulating Aβ into the brain [17–19]. The balance between LRP1-mediated efflux and RAGE-mediated influx is therefore crucial in regulating brain Aβ levels [20]. A mathematical representation of the BBB that captures the physical behavior of the receptor-based boundary is essential for reliable simulation of Aβ clearance from the brain. Currently, there is no available mathematical model in the literature that accounts for the size, number, and kinetics of receptors on both sides of the BBB.

A foundational framework for modeling receptor-mediated diffusion processes was laid by Berg and Purcell [21], who developed a theory for the accuracy of chemoreception in biological systems by considering the diffusive flux of particles to a spherical surface sparsely decorated with receptors. They demonstrated that the flux of ligands to receptors scales linearly with both the receptor size and number, and they introduced the concept of a “diffusive limit” for sensing, which remains central in quantitative models of cell signaling. This classical model was extended by introducing time-dependent treatments of ligand binding to spheres partially covered by receptors [22]. This model showed that the early-time binding kinetics differ significantly from steady-state assumptions and that the total flux approaches the Berg–Purcell prediction only after a transient phase. A central aspect of many receptor-based models is the incorporation of Michaelis–Menten kinetics, originally developed to describe enzyme-catalyzed reactions [23]. This kinetic framework assumes that the reaction rate is proportional to the concentration of the ligand and saturates as receptor sites become occupied. In the context of diffusive transport and boundary models, the Michaelis–Menten formalism appears as a non-linear Robin boundary condition that captures saturable binding at the surface. Specifically, the flux into the boundary is modeled as 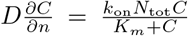, where *C* is the concentration, *n* is the normal vector to the surface, *k*_on_ is the binding rate constant, *N*_tot_ is the total receptor density, and *K*_*m*_ is the Michaelis constant. This form was explored by considering dynamic traps that switch between active and inactive states, effectively implementing a time-dependent Michaelis–Menten-type behavior at the boundary [24].

Further refinements involved homogenization techniques where a patchy absorbing surface could be approximated using a Robin boundary condition—an approach which replaces a mixed Dirichlet–Neumann boundary with an effective single boundary of the form 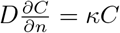, where *κ* encodes partial absorption [25, 26]. This effective medium theory simplifies modeling of surfaces with distributed receptors without resolving each patch individually. It has been formalized for narrow-escape and mean first-passage time problems on spherical targets with small absorbing patches [27], and extended to planar surfaces with periodic patches [28]. Further developments incorporate anisotropic diffusion and rotational effects for patchy particles [29]. More recently, the classical Berg–Purcell model was revised by including finite receptor kinetics, showing that traditional models can greatly overestimate uptake rates [30].

In this current study, we extend the receptor-mediated diffusion model with finite receptor kinetics [30] to develop a boundary condition for the BBB. To achieve this, we update the previous model to account for two domains – brain and blood – with the BBB at their interface. This is the first BBB boundary model that incorporates the number, size, and kinetics of BBB receptors. We couple this boundary condition with a diffusion model for Aβ transport through the brain. Aβ monomers pass between the two domains via receptors on each side, while maintaining mass conservation. First, we derive the boundary condition using brain-specific and Aβ-specific assumptions. Then, we solve the steady-state diffusion problem analytically using the derived boundary condition. We next solve the problem numerically to analyze BBB behavior in brain waste clearance; we both verify our numerical method against analytical results and validate the predicted Aβ concentrations against measurements reported in the literature. We highlight the sensitive dependence of Aβ on a few key parameters which are known to vary with aging.

Finally, we adapt our numerical simulations to investigate transient Aβ changes in one complex physiological and one complex pathological scenario. Sleep has been linked to decrease in Aβ concentration in the brain [31], and growing evidence suggests sleep disruption contributes to Alzheimer’s pathology [32, 33]. We perform sleep simulations that indicate rapid eye movement (REM) sleep may play a vital role in Aβ clearance. The complex pathological scenario we consider is spreading depolarization, which is a wave of abnormal neuronal activity that propagates through the cerebral cortex and occurs due to a loss of brain ion homeostasis [34]. Spreading depolarization (SD) is associated with migraine with aura (as well as many other neurological disorders) [35], and migraine is linked to an increased risk of dementia [36, 37]. We perform simulations that predict transient changes in Aβ concentration due to SD which may help explain such associations.

## II. METHODS

### A. Boundary Homogenization

We start by creating a general receptor-mediated diffusion problem involving two domains, as illustrated in Figure 1. In this model, diffusing substrates (with concentration *C*_*α*_, *α* = {*i, o*}) can pass through the distributed transporters on the boundary Γ separating the inner (Ω_*i*_) and outer (Ω_*o*_) domains. These substrates can bind to enzymes (*E*_*α*_) on the walls, which act as transporters, allowing the substrates to travel between the two domains. During this process, a substrate that binds to an enzyme forms a bound complex (*B*_*α*_). The enzymes responsible for transporting substrates from the outer to the inner domain may differ from those facilitating transport from the inner to the outer domain. Once a bound complex is formed, the substrate can either be transported across the domain, or released back into its starting domain. Additionally, a bound complex can only transport one substrate at a time.

**FIG. 1.**
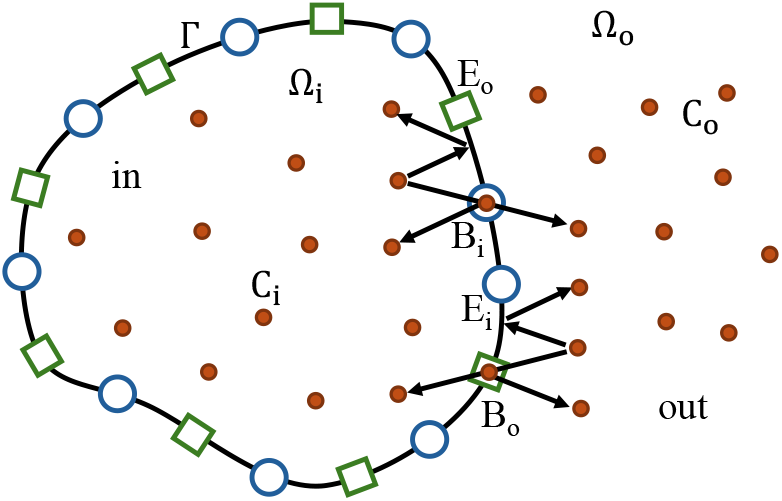
Schematic of solute transport between two domains via binding receptors. Here, the two “in” and “out” domains are separated by the surface Γ, and the inner (*C*_*i*_) and outer (*C*_*o*_) substrates can be transported across the boundary between the domains. *E*_*α*_ and *B*_*α*_ represent the enzyme and bound complex for the inner and outer receptors, respectively.

The advection-diffusion equations for the inner and outer domains (indicated with subscripts *i* and *o*, respectively) can be written as follows

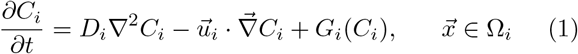

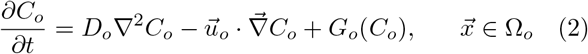

where *C*_*α*_ is a scalar field representing the concentration of the substrate, *D*_*α*_ is the substrate’s diffusivity,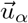 represents the velocity field, and *G*_*α*_ accounts for substrate generation within the domain. To solve the system of Equations (1) and (2), we require a boundary condition for the wall that separates the domains which is defined as

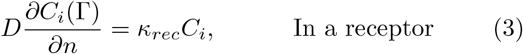

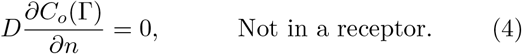

To homogenize the boundary condition in Equation (3) we replace it with a homogeneous boundary condition as follows

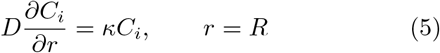

where trapping rate *κ* is chosen to incorporate the number, size, and arrangement of receptors for the specific system. We apply the classical substrate and enzyme reaction scheme at the boundary for both domains, as follows [38]:

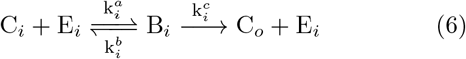

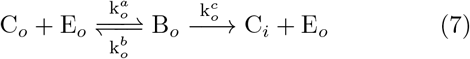

where *k*^*a*^, *k*^*b*^, and *k*^*c*^ represent the rates of association, breakup, and catalysis, respectively. Initially we assume that these rates are different for receptors associated with each direction. Additionally, since the receptors for each direction are different, the enzyme *E*_*α*_ and the bound complex *B*_*α*_ are not equal. Using chemical Equations (6) and (7) and the law of mass action, we can formulate a homogeneous boundary condition for the flux exiting the inner domain as

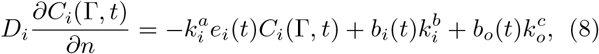

where *e*_*i*_(*t*) and *b*_*i*_(*t*) are free receptor and bound receptor concentrations respectively and satisfy the following equations

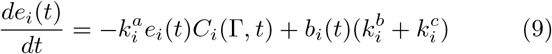

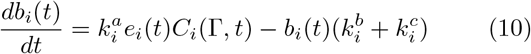

which satisfies the total receptor concentration. From Equations (9) and (10), one can obtain the conservation equation *e*_*i*_(*t*) = *e*_0,*i*_ − *b*_*i*_(*t*), where *e*_0,*i*_ represents the number of receptors per unit surface area. Consequently, we can eliminate *e*_*i*_(*t*) from Equations (8) and (10), rewriting them as:

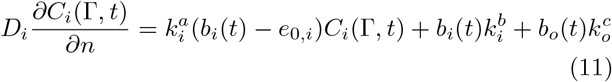

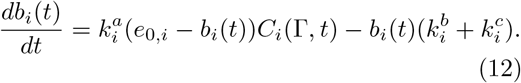

In cases where the volume concentrations of receptors are considerably smaller than those of the substrates, Briggs and Haldane’s derivation adopts the steady-state approximation [39]. This approximation is based on the observation that in most systems (including the BBB system that is the focus of this work), the concentration of the bound complex will quickly reach a steady state after an initial burst phase, then its concentration will remain relatively stable until a significant amount of substrate has been consumed. Therefore, we can rewrite Equation (12) using the steady-state approximation (*db/dt* = 0) as follows:

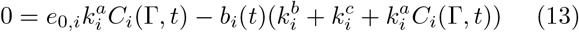

which leads to:

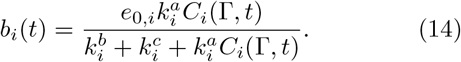

By substituting this expression into Equation (11), we have:

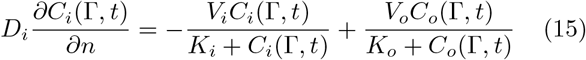

where the maximal reaction velocity and half-saturation constant for each domain are defined as follows:

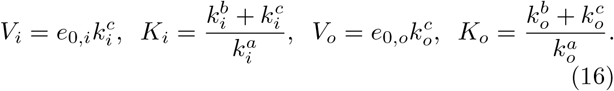

Following the same steps for the flux entering the outer domain yields the same flux but with a different sign, ensuring mass conservation,

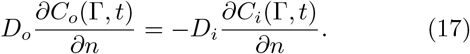

This completes the derivation of the boundary condition for the membrane separating the two domains in the problem introduced in Equations (1) and (2).

### B. Aβ transport across the BBB

Aβ is a waste protein that is naturally generated in the brain and must be effectively removed. Failure to clear Aβ from the brain leads to its accumulation in the interstitial space, contributing to the development of diseases such as Alzheimer’s [3]. Approximately half of the secreted Aβ monomers are removed through the BBB and drain directly into the bloodstream [5, 10]. This transport occurs via active transporters in the endothelial cells lining the vessel walls, which bind to the monomers and transfer them across the barrier [12, 13].

We adapt the introduced boundary condition to simulate Aβ transport through the BBB. The domain consists of two concentric cylinders, where the inner cylinder represents the capillary, and the outer cylinder represents the surrounding brain tissue. Figure 2 illustrates the concentration distribution for a capillary surrounded by brain tissue, where transport occurs. Equations (1) and

**FIG. 2.**
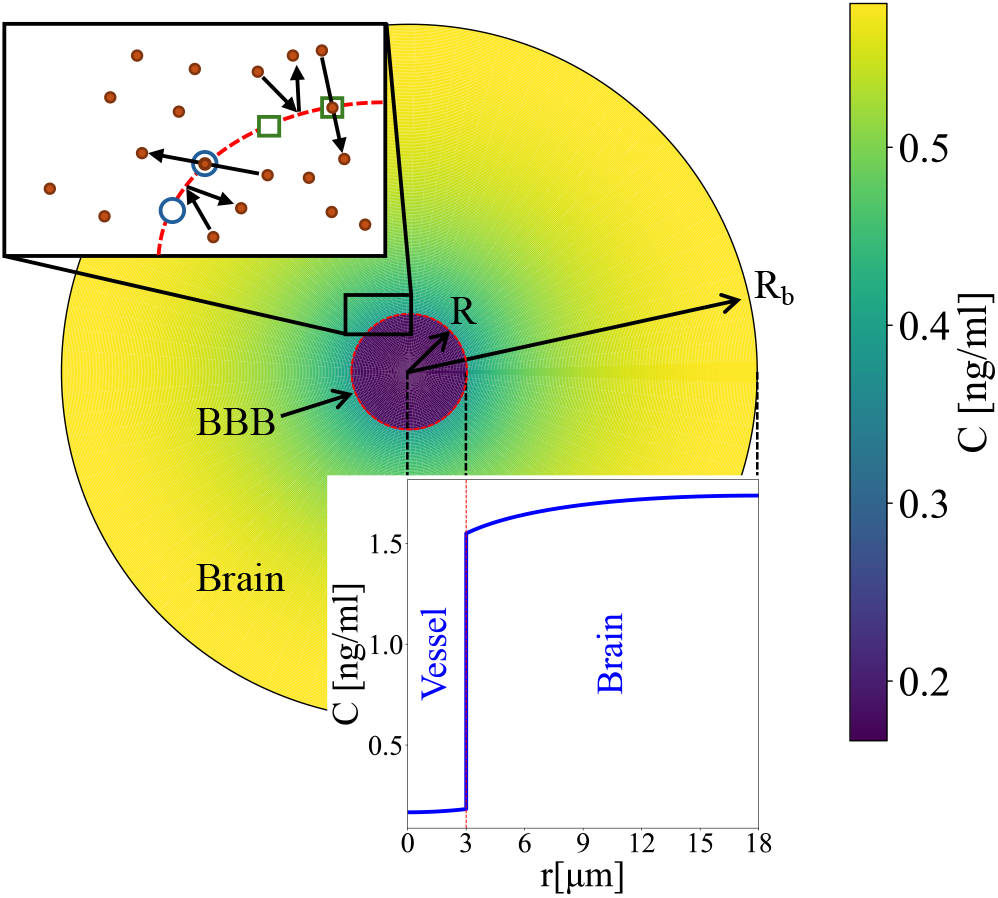
A schematic diagram of the axisymmetric simulation domain. The color contour indicates the Aβ concentration *C*. The one-dimensional plot (bottom) quantifies the concentration profile as a function of radius *r*. The zoomed view of the boundary (top left) illustrates the BBB receptor kinetics that we model.

(2) can be modified to capture this transport by considering waste removal in the vessel due to blood advection (inner domain) and the natural generation of Aβ in the brain (outer domain). The modified equations, under the assumption of axisymmetry, are as follows:

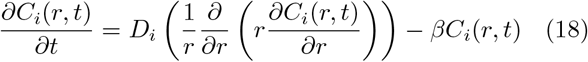

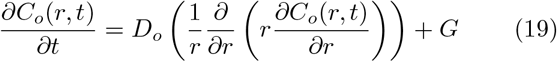

where *β* is blood sink coefficients, a constant accounting for Aβ clearance in the blood due to advection and *G* is the production of Aβ that occurs naturally in the brain. Beyond the shared boundary conditions at the BBB, one additional boundary condition is required for each domain. The following boundary conditions are assumed for the inner and outer domains due to the symmetry of the geometry:

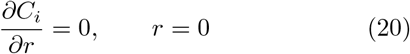

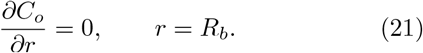

The boundary condition introduced in Equation (15) is adjusted to represent Aβ transport between the two domains. We assume *k*^*b*^ = 0 in Equations (6) and (7), meaning that when Aβ monomers bind to the BBB receptors, they are transferred to the other side without dissociation. The trapping rate *κ* in Equation (5) is defined as

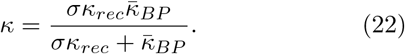

In this expression *κ*_*rec*_ is the rate that a receptor binds a substrate. Additionally, surface trapping rate 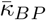 and receptor area fraction *σ* are defined as follows

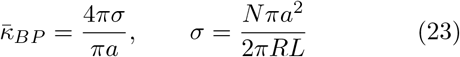

where *a* and *N* represent the receptor radius and the number of receptors, respectively. By choosing *ϵ* as the ratio of the receptor radius to the vessel radius and substituting Equation (23) into Equation (22), we obtain

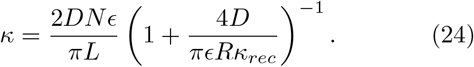

Taking *κ*_*rec*_ *→ ∞* [30] one can plug *κ* into Equation (5). Additionally, in the case of free receptors, one can assume *b*(*t*) = 0, and Equation (11) reduces to Equation (5). Therefore, assuming a similar radius for the receptors that transfer Aβ from brain to blood as for those that transfer from blood to brain (*ϵ*_*i*_ = *ϵ*_*o*_), and assuming a small thickness of the vessel wall, the rate of association for the blood and brain sides becomes

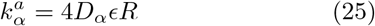

where *α* = {*i, o*}. The receptor concentration per unit surface area, *e*_0_, for a cylindrical vessel can be calculated as

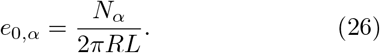

Additionally, we assume that the rate of catalysis (*k*^*c*^) is the same for both the blood and brain sides. Applying these assumptions to Equation (16), Equation (15) can be rewritten as follows:

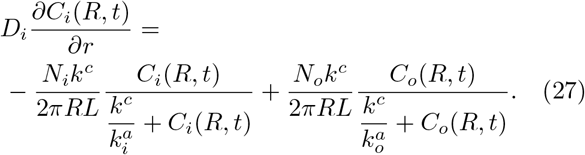

The nondimensional form of this boundary condition, as well as the governing equations (18-19), are available in §I of the Supplemental Materials [40]. The complexity of this boundary condition arises from the fact that it accounts for the number, size, and kinetics of the BBB receptors, as well as the saturation of transport due to high concentrations at the boundary. To assess the significance of these complexities, we performed simulations using a simplified linear form of the boundary condition and observed a significant difference compared to the results obtained with the complete version. This comparison is provided in §II of the Supplemental Materials [40].

### C. Numerical Method

As mentioned, we consider an averaged cross-sectional domain consisting of a capillary and its corresponding surrounding portion of brain tissue (Figure 2). In this configuration, variations in axial direction are averaged, and the problem exhibits azimuthal symmetry. Therefore, the model is reduced to an axisymmetric one-dimensional (1D) cylindrical problem with variations in the *r*-direction, referenced from the center of the capillary. To solve the problem with the introduced boundary conditions, we developed a numerical model using the Finite Difference Method. We employed an explicit time integration scheme with the forward Euler method, which is second-order accurate in space (based on central finite differences) and first-order accurate in time. The implementation details of the boundary condition in Equation (27) are provided in §III of the Supplemental Materials [40].

First, we nondimensionalized the equations and boundary conditions and simulated the problem for both transient and steady-state cases (see §I in the Supplemental Materials [40]). The vessel and brain domains are simulated within separate computational domains, and the interface boundary condition is handled by applying the flux introduced in Equation (27) at the right boundary of the vessel domain and the left boundary of the brain domain. To ensure steady-state, a criterion of 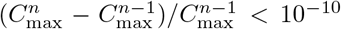 is enforced, where *n* represents the iteration number and *C*_max_ is the highest concentration in the domain. Figure 2 shows a contour and a profile of concentration distribution in the vessel and brain.

### D. Parameter Range

To perform simulations, we need to specify the parameters used in the equations. Since determining some biological values requires challenging in vivo experimental measurements we focus on mice brain where more data is available and as these measurements are prone to substantial biological variability and large uncertainties, we test a range of these parameters. The radius of capillaries ranges from 2.5 µm, approximately the size of a red blood cell, to 4 µm. Additionally, as mentioned, LRP1 is primarily responsible for mediating the efflux of Aβ from the brain interstitial fluid into the blood and RAGE is involved in transporting circulating Aβ from the blood into the brain. In Equation (27), we specify the number of these receptors per unit length (*N/L*), and the *N*_*o*_*/N*_*i*_ ratio determines the weight of transport in each direction.

The diffusivity of Aβ in a free medium is *D* = 2.3 *×* 10^−10^ m^2^/s [41]. This value is used for Aβ diffusion in the blood (*D*_*i*_ = *D*). To account for the porous nature of brain tissue, the diffusivity of Aβ is modified by the tortuosity of the brain tissue, *D*_*o*_ = *D*_*eff*_ = *D/λ*^2^ where *λ* = 2.04 [41]. Additionally, the size of the domain (*R*_*b*_) is defined as half of the average distance between capillaries in mice brain, which is approximately 36 µm [42]. A list of parameter ranges used for the simulation is provided in Table I. A column of default values is provided in the table, and unless otherwise stated in the figure captions, all results use these default values.

**TABLE I.** Parameters used in simulations.

| Parameter | Definition | Range | Default value | Reference |
| --- | --- | --- | --- | --- |
| $N_i/L$ | Number of LRP1 receptors | $[2 \times 10^3, 2 \times 10^4]$ #/$\mu\text{m}$ | $2 \times 10^4$ #/$\mu\text{m}$ | Kyrtsos et al. [43] |
| $N_o/L$ | Number of RAGE receptors | $[2 \times 10^3, 2 \times 10^4]$ #/$\mu\text{m}$ | $2 \times 10^4$ #/$\mu\text{m}$ | - |
| $a$ | Receptor radius | | $10^{-3}$ $\mu\text{m}$ | - |
| $\beta$ | Blood Sink Coefficient | $[1, 10^3]$ $\text{s}^{-1}$ | $10$ $\text{s}^{-1}$ | - |
| $k_c$ | Turnover rates of transporter | $[1, \infty]$ $\text{s}^{-1}$ | $10$ $\text{s}^{-1}$ | - |
| $R$ | Capillary radius | $[2.5, 4]$ $\mu\text{m}$ | $3$ $\mu\text{m}$ | Miyawaki et al. [44] |
| $R_b$ | Capillary-capillary distance | | $36$ $\mu\text{m}$ | Yousofvand et al. [42] |
| $D$ | Aβ diffusivity | | $230$ $\mu\text{m}^2/\text{s}$ | Sykova et al. [41] |
| $\lambda$ | Tortuosity of brain | | $2.04$ | Sykova et al. [41] |
| $G$ | Aβ production | | $0.1$ $\text{ng}/(\text{ml}\cdot\text{s})$ | Yousofvand et al. [42] |

The introduced LRP1 receptor number is used as the default value for both LRP1 (*N*_*o*_*/L*) and RAGE (*N*_*i*_*/L*), and variations in *N*_*o*_*/N*_*i*_ relative to this default value are analyzed. Although a range of capillary sizes is considered, a default size of 3.3 µm is used for simulations. Additionally, since half of the generated Aβ monomers are removed by the BBB, we consider half of the production rate provided in Table I for the source term *G* in Equation (19) (i.e., we exclude the half of production that would be cleared via CSF).

## III. RESULTS

### A. Steady-state results

After creating the domain and defining the boundary conditions, we perform simulations for various cases. We begin with steady-state analysis of the concentration profile in the domain for different blood sink coefficients (*β*) and receptor number ratios (*N*_*o*_*/N*_*i*_). Figure 3(a) illustrates the concentration profile for different receptor number ratios with *β* = 10 s^−1^. The saturation level is highly dependent on the receptor number ratio, with higher saturation concentration observed on the side of the boundary with fewer receptors. Interestingly, we find that the concentration profile inside the vessel is independent of receptor number ratios and depends only on *β*. From the analytical solution for the steady-state concentration inside the vessel, provided in §IV of the Supplemental Materials [40], we observe that the concentration in the vessel is in fact independent of the receptor number ratio, which verifies our numerical model.

**FIG. 3.**
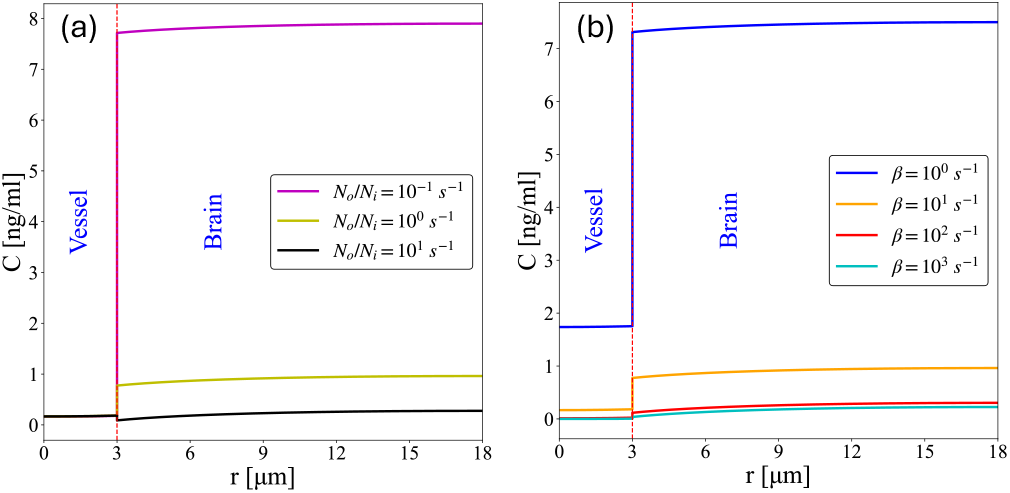
Steady-state concentration profiles for (a) different *β* values with *N*_*o*_*/N*_*i*_ = 1 and (b) different *N*_*o*_*/N*_*i*_ with *β* = 10 *s*^−1^.

An important observation in Figure 3 is the concentration level of Aβ calculated in the vessel. It has been shown that the concentration of Aβ in plasma in the mouse brain is approximately 0.1-0.2 ng/ml [45]. A similar concentration level is observed in our simulations for *β* = 10 *s*^−1^. This agreement is achieved using the boundary condition from Equation (27), along with the receptor numbers and the production rate of Aβ specific to the mouse brain, thereby validating our model.

Figure 3(b) shows the concentration profile for different *β* values with *N*_*o*_*/N*_*i*_ = 1. A higher *β* corresponds to stronger clearance from the blood, leading to a lower concentration in both inner and outer domains. The shape of the profile varies for different *β* values and part of that is due to the different diffusivity between the two domains. For higher *β* values, the concentration in the vessel decreases, approaching zero. Beyond this point, the vessel concentration can no longer drop further, but increasing *β* still leads to enhanced clearance, resulting in a smaller concentration difference between the vessel and the brain.

To gain a better understanding of the effects of *β* and *N*_*o*_*/N*_*i*_ = 1 on the concentration in the blood and brain, we performed simulations across a wider range of these parameters. The averaged steady-state concentrations in the vessel and the brain are shown in Figure 4 for different receptor number ratios and *β* values. As seen in Figure 4(a), increasing the number of receptors on the brain side compared to the blood side leads to a lower average concentration in the brain, as more receptors facilitate the transport of Aβ from the brain to the blood. This behavior is not observed for the concentration in the vessel, where the average concentration remains independent of receptor number ratios (Figure 4(b)), the same principle we observed in Figure 3(a).

**FIG. 4.**
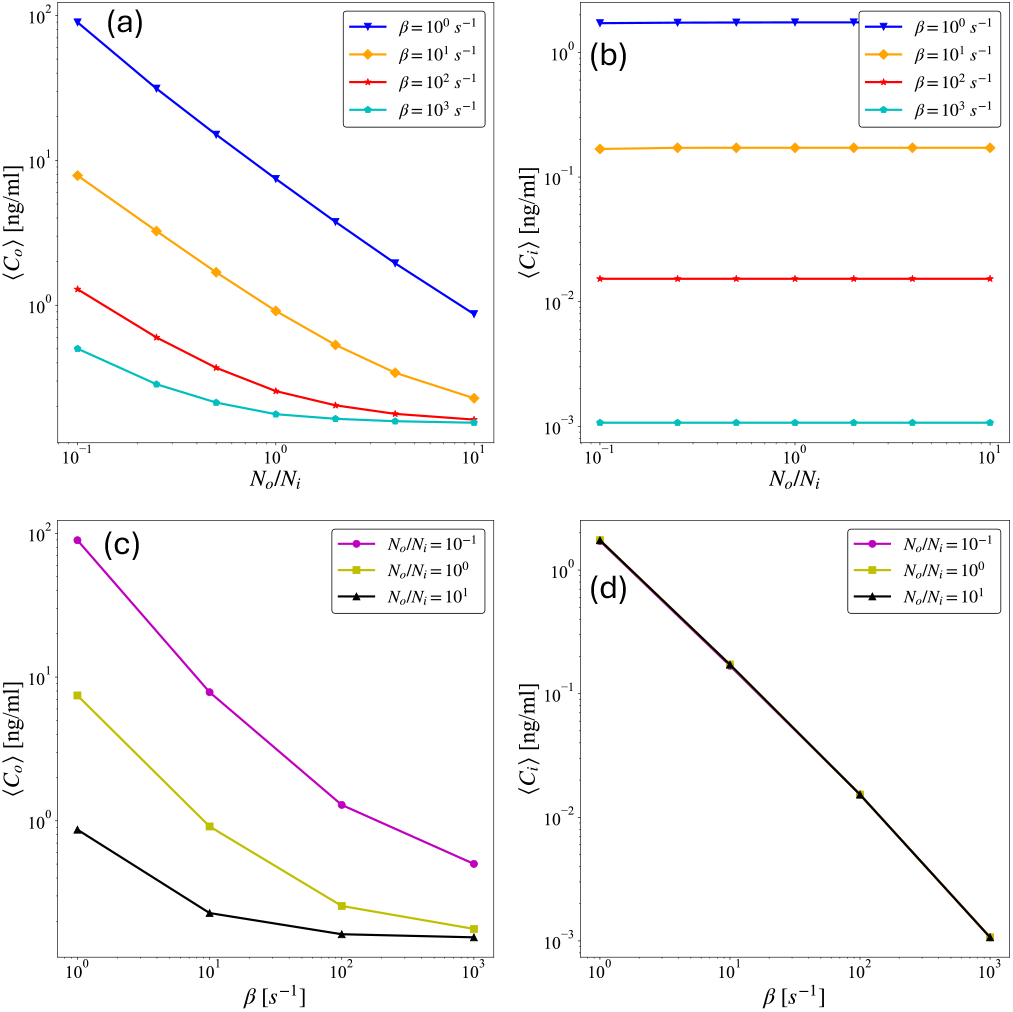
Spatially-averaged steady-state concentrations for different *β* and *N*_*o*_*/N*_*i*_. (a-b) Average concentration as a function of *N*_*o*_*/N*_*i*_ for the (a) inner (vessel) or (b) outer (brain) domain. (c-d) Average concentration as a function of *β* for the (c) inner (vessel) or (d) outer (brain) domain.

Figures 4(c) and 4(d) show the variation of concentration in the brain and vessel with *β*. Since *β* represents the strength of clearance by advection in the vessel, increasing *β* results in a decrease in concentration in both domains. However, this dependency is more prominent in the vessel, as the interface boundary acts as a barrier between the concentration in the brain and the sink in the blood.

Another important parameter we analyzed is the vessel size, which can affect the BBB and the concentration of Aβ in the brain. Figures 5(a) and (b) show the concentration profiles and the variation of averaged concentration for different vessel diameters, respectively. Here, we vary only the vessel diameter while keeping all other parameters constant, including the receptor-to-vessel radius ratio (*ϵ*) and the total number of receptors. Since there is a sink term in the vessel, an increase in vessel diameter results in a stronger sink effect, leading to lower concentrations in both the vessel and the brain. As the concentration decreases, the level of saturation in the brain tissue also decreases with increasing vessel size.

**FIG. 5.**
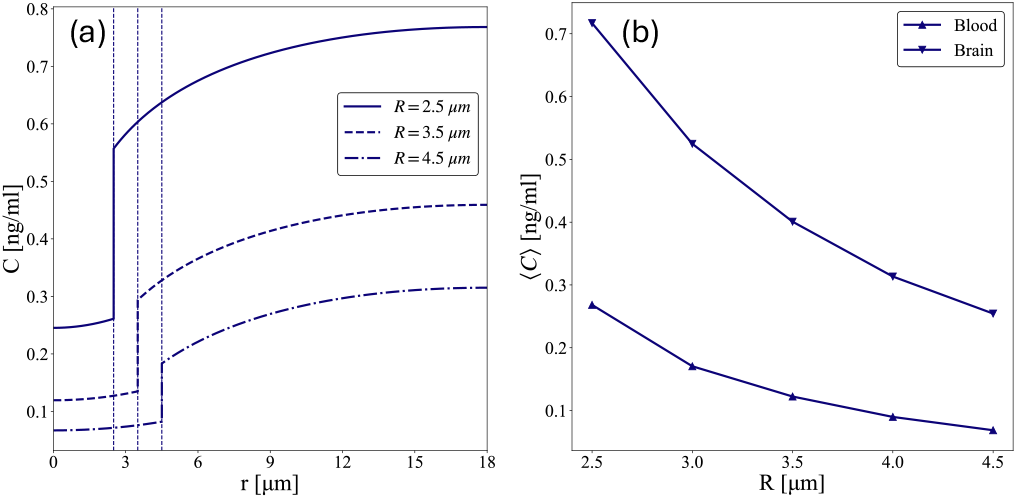
Steady-state concentration varies with vessel radius. (a) Concentration profiles for three different capillary radii. (b) Spatially-averaged concentration for the inner (vessel) and outer (brain) domains as a function of capillary radius.

### B. Transient results

So far, we have analyzed the effect of different parameters on the transport rate of Aβ through the BBB in steady state. Here, we present results at different time instants to analyze the transient behavior of the system under various brain dynamics. We begin by simulating the transient behavior of Aβ concentration levels in the brain in response to changes in blood flow. First, we perform a steady-state simulation corresponding to the case of *β* = 1 s^−1^ and *N*_*o*_*/N*_*i*_ = 2 and use these results as the initial condition. Then, the transient simulation begins by using this initial condition and increasing *β* to 10 s^−1^ until reaching a new steady state. The same criterion of 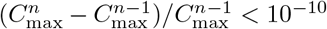 is enforced for identifying a steady state. Figure 6 shows the concentration pro-file and the average concentration in the vessel and brain for these transient simulations. Panels (a) and (b) in Figure 6 illustrate the evolution of concentration profiles and averaged concentration respectively, and the time scale of these changes. This analysis demonstrates the effect of increasing blood flow speed on Aβ clearance from the brain. The time required to reach the new steady state when increasing *β* from 1 to 10 s^−1^ is approximately 1 min.

**FIG. 6.**
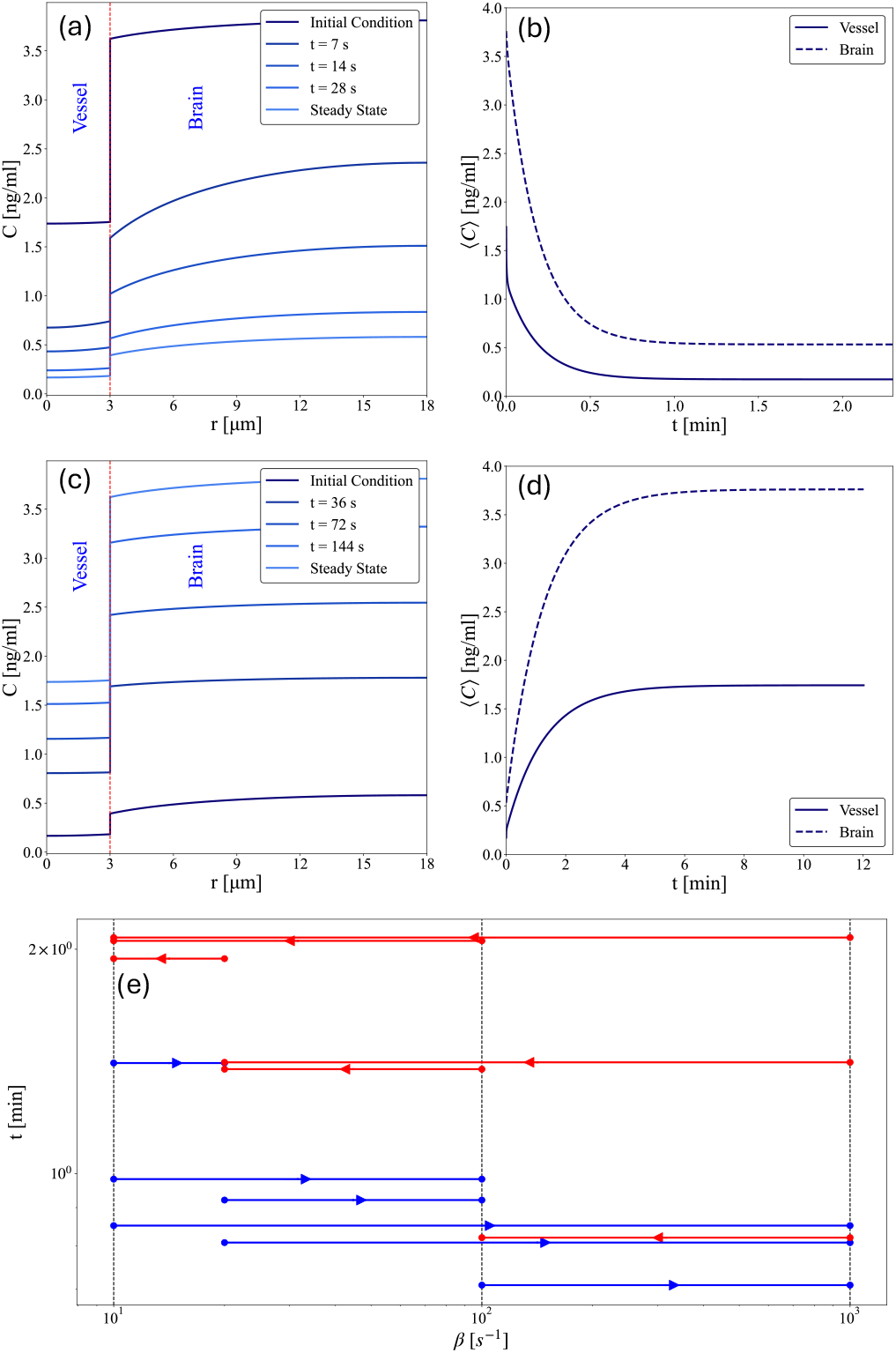
Transient times are shorter for increasing *β* than decreasing *β*. (a) Transient variation of the concentration profile and (b) spatially-averaged concentration when *β* is increased from 1 to 10 s^−1^. (c) Transient variation of the concentration profile and (d) spatially-averaged concentration when *β* is decreased from 10 to 1 s^−1^. (e) Time needed to reach steady state for different increases and decreases of *β*. All plots correspond to *N*_*o*_*/N*_*i*_ = 2.

We next performed the reverse analysis: we take the steady-state results of the case with *β* = 10 s^−1^ as the initial condition and begin the transient simulation with *β* equal to 1 s^−1^ (Figures 6[c-d]). This analysis characterizes to the effect of reducing blood flow on brain Aβ concentration. An interesting observation here is that the time required to reach steady state is more than five times longer than the previous case. This indicates that increasing blood flow reduces the concentration in the brain much faster than decreasing blood flow increases the concentration. We performed a similar analysis for various 2- 10- or 100-fold increases and decreases in *β* and recorded the time required to reach steady state (Figure 6[e]). All the red cases, which represent a *decreased* value of *β*, consistently take longer to reach steady state compared to the analogous increased *β* (blue) cases. Additionally, due to the complex nature of the BBB boundary condition, no clear trend can be identified within the blue (increase) or red (decrease) cases when considered separately. These comparisons strongly motivate numerical investigation of phenomena that alter cerebral blood flow and/or capillary diameter, such as sleep and spreading depolarization.

To analyze the dynamics of Aβ concentration in the brain and blood during sleep, we simulated a sleep cycle for mice. We assumed a cycle consisting of 8 min of non-REM (NREM) sleep and 1 min of rapid eye movement (REM) sleep [46]. While it has been shown that arterial diameter changes during different sleep phases [47], the size of capillaries remains constant. However, the velocity of blood flow during REM is approximately twice as high as during NREM and awake states [48]. We simulated a transient scenario for REM and NREM phases using *N*_*o*_*/N*_*i*_ = 2 and varied *β* between 10 s^−1^ and 20 s^−1^. The results are shown in Figure 7. In each REM cycle, the average concentration of Aβ in the blood vessel and the brain decrease by 51% and 34%, respectively,and equilibrium at that lower concentration is reached after about half the REM duration. This suggests that REM sleep may, in addition to a myriad of other vital functions [49], aid the brain in removing Aβ. Note that the difference in transient time (required to reach steady state) in this case is substantially less than that of Figure 6(c-d) because the change in *β* is smaller.

**FIG. 7.**
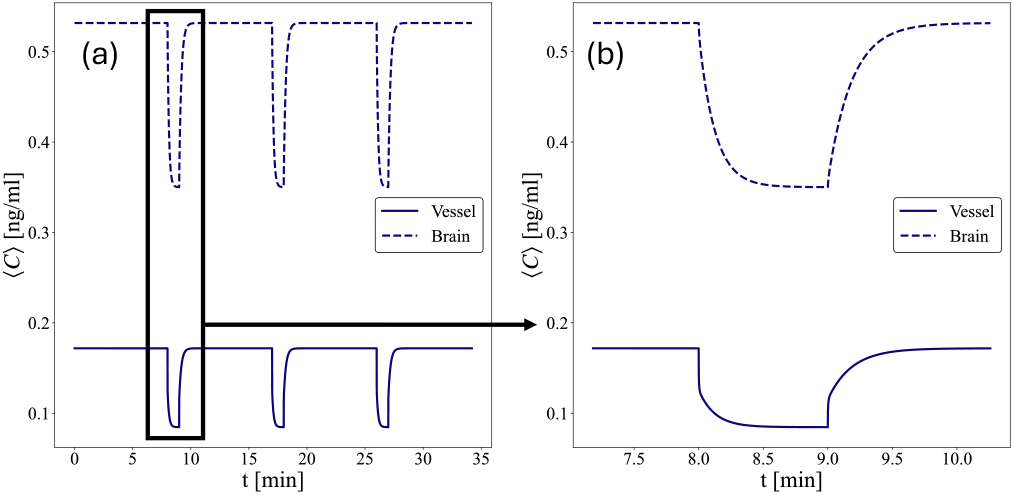
Aβ concentration during sleep suggests REM sleep may play an important role in waste clearance. (a) Three sleep cycles consisting of 8 min of NREM sleep and 1 min of REM sleep. (b) A zoomed-in enlargment of the 2.5-min period indicated in (a). In all cases, *β* changes between 10 and 20 s^−1^ in NREM and REM sleep, respectively.

Finally, we simulated transient effects due to a pathological phenomenon known as “spreading depolarization” (SD). As mentioned above, SD occurs due to loss of brain ion homeostasis that propagates as a wave through the cerebral cortex [34]. During this process, the size of the brain vasculature varies over time, with capillaries experiencing three sequential phases. Phase I lasts for 1 min and involves vasoconstriction, with an average decrease of 8.2% in capillary radius compared to baseline (this average is taken across first-, second-, and third-order capillaries). Phase II follows Phase I and lasts for about 3 min, during which the capillary radius increases by an average of 12.5% relative to baseline. Finally, in Phase III, a persistent vasoconstriction occurs, reducing the capillary radius by 11.2% compared to baseline [50]. We performed a time-varying simulation of this process using default values of *β* = 10 s^−1^, *N*_*o*_*/N*_*i*_ = 1, and a baseline capillary radius of 3 µm. As shown in Figure 8, the concentration of Aβ varies in response to changes in capillary size, and the different timescales of these phases lead to a reduced then increased steady-state Aβ concentration during Phases II and III, respectively. In the latter case, Aβ is elevated by 28% and 22% (compared to the baseline concentration before SD) in the vessel and brain tissue, respectively. These transient simulations demonstrate the capability of the derived BBB boundary condition to model different brain states and the dynamic behavior of brain vasculature.

**FIG. 8.**
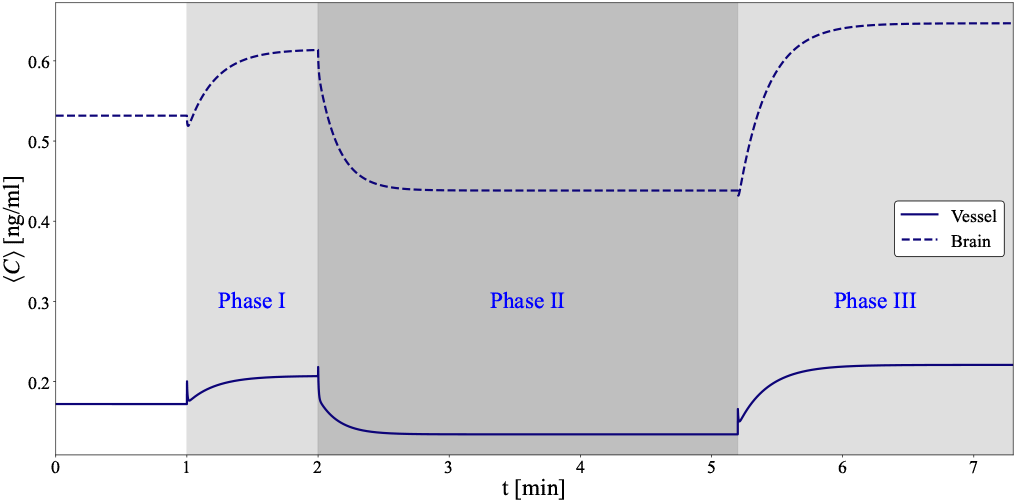
Averaged Aβ concentration variation in the brain and vessel for different phases of spreading depolarization for mice brain. Phase I: 60 seconds, 8.2% decrease in capillary radius relative to baseline, Phase II: 3 minutes, 12.5% increase in capillary radius relative to baseline, Phase III: persistent, 11.2% decrease in capillary radius relative to baseline. Note that the brief jumps or drops in the concentrations at the beginning of each phase are small numerical artifacts that arise due to the concentration profile adjusting following sudden changes in the size of the vessel/brain domain.

## IV. CONCLUSION

In this study, we developed a novel mathematical model to describe the blood-brain barrier (BBB), incorporating receptor kinetics and accounting for both the number and properties of active transporters on either side of the BBB. Starting from a finite receptor kinetics model, we extended boundary homogenization to derive an effective boundary condition for solute flux between two domains representing the brain and the blood. This new boundary condition accounts for bi-directional transport between the two domains ensuring mass conservation, and it incorporates nonlinear receptor-mediated kinetics, enabling the first mathematical representation of BBB transport processes. We used this boundary model to simulate Aβ transport between the brain and blood, largely motivated by the important role that impaired Aβ clearance plays in pathogenesis of Alzheimer’s disease and related dementias [2, 51]. We applied the derived boundary condition to an axisymmetric cylindrical geometry, representing a capillary surrounded by brain tissue, and incorporated relevant biophysical parameters such as receptor density, diffusivity, and Aβ production rates. The governing equations were nondimensionalized (Supplementary Material §I) and solved numerically using a finite difference method. Both steady-state and transient simulations were performed to investigate the influence of key parameters such as receptor number ratio, blood clearance rate, and vessel diameter on Aβ distribution. Our numerical results were verified against the steady-state analytical solution (Supplementary Material §IV) and then leveraged to characterize parametric dependencies.

Our results show that the receptor number ratio *N*_*o*_*/N*_*i*_ significantly affects Aβ concentration in the brain but not in the blood, consistent with the analytical solution. This sensitivity to receptor number ratio may, in part, explain why aging is the greatest risk factor for Alzheimer’s disease [52]. A prior study reported that LRP1 expression *decreases* up to 6.4-fold and RAGE *increases* up to 82-fold with aging (in mice) [53]. Hence, *N*_*o*_*/N*_*i*_ decreases by over two orders of magnitude across the murine lifespan, suggesting that the brain Aβ equilibrium concentration could increase 10-fold or more (Figure 4c) due to this factor alone.

The blood clearance rate *β* was also found to play a very significant role in regulating Aβ levels in both compartments, with increasing *β* lowering concentrations more rapidly in the vessel than in the brain. Specifically, increasing *β* by a factor of 1000 decreased blood and brain Aβ concentration by about three orders of magnitude and up to about two orders of magnitude, respectively (Figure 4c-d). Notably, the change in Aβ concentration in the blood is independent of *N*_*o*_*/N*_*i*_, but the *rate of change* in brain concentration with the respect to *β* is greater when *N*_*o*_*/N*_*i*_ is smaller, as occurs with aging (compare purple and black curves in Figure 4c). This leads to an interesting observation with important therapeutic implications: although aging-related decrease in *N*_*o*_*/N*_*i*_ likely leads to larger Aβ concentrations in brain tissue, our model suggests there is an increasing sensitivity to elevated blood flow in this regime. Hence, approaches for elevating cerebral blood flow, such as exercise [54] or neuromodulation [55], may be especially important for reducing A*β* load in old age. A related prediction from our model involves the temporal asymmetry that arises in transient simulations: increasing blood flow led to faster reduction in brain Aβ than the rate of Aβ increase observed when blood flow was reduced (Figure 6). Hence, even brief increases in cerebral blood flow (such as during REM sleep) may have a disproportionately large effect on Aβ clearance. Our model also predicts that reductions in blood flow (i.e., decreased *β*) in aging (i.e., when *N*_*o*_*/N*_*i*_ decreases) will lead to substantial increases in brain Aβ concentrations. There is evidence suggesting this effect contributes to pathogenesis of cognitive impairment and dementia, although causality has been difficult to establish in experiments [56].

We demonstrated our model’s applicability to complex physiology by simulating sleep as a cycle of variations in blood flow (captured by varying *β*). Based on a two-fold change in blood flow, motivated by published experimental measurements [48], our model predicts that the Aβ concentration in the blood and brain each decrease by 50% and 34% (during the REM phase, compared to NREM), respectively. An important, open question concerns the evolutionary necessity of two-phase (NREM and REM) sleep across most animal species [57]. Our results (Figure 7) suggest that REM sleep may play an important role in enhancing Aβ clearance. Indeed, the time scale for murine REM sleep (about 1 min) is only a bit longer than the time required to reach equilibrium at the lower Aβ concentration. This can be further rationalized by considering another waste clearance pathway in the brain, the glymphatic system, which is most active during NREM sleep [58]. The glymphatic system is an important pathway for solute exchange between cerebrospinal fluid (CSF) and interstitial fluid [3]. We have recently shown that Aβ clearance depends on both BBB and glymphatic function, and that overall clearance is synergistically amplified when both clearance mechanisms are present [42]. Hence, the glymphatic system may transport interstitial Aβ to the vicinity of blood vessels during NREM sleep and BBB efflux – combined with elevated blood flow – may enhance Aβ clearance during REM sleep. In future experimental and numerical studies, we intend to test this hypothesis about how the structure of a sleep cycle may maximize brain Aβ clearance.

We also simulated a complex pathological phenomenon, spreading depolarization (SD). SD refers to the loss of brain ion homeostasis, which propagates through the cortex as a reaction-diffusion wave and is associated with migraine with aura, stroke, traumatic brain injury, and more [34]. SD elicits changes in capillary diameter, likely due to SD-related calcium ionic signaling to pericytes (cells that wrap around capillaries) [50]. Our model suggests that changes in capillary diameter alone can lead to large disruptions in brain Aβ concentration (Figure 8), with decreases as much as 18% for more than 2 min (Phase II) and increases by up to 22% for over 1 min (Phase III). This highlights the prospect of ionic manipulation of pericytes to augment brain microcirculation [59], potentially increasing Aβ clearance. A limitation of our approach is that we only modeled the variation in vessel diameter and not the associated complex changes in blood flow, including flow stagnation, which has been reported in prior studies [60, 61]. When this additional factor are accounted for, the maximum increase in brain Aβ is likely to be significantly higher that our simulations predict. These elevated Aβ concentrations may help explain why migraine has been associated with an increased risk of dementia [36, 37].

There are several notable limitations of this study. Our model geometry is highly idealized, consisting of a 1D (i.e., axisymmetric two-dimensional) domain. While this assumption facilitates ease of data interpretation and computational tractability, true brain vasculature is much more heterogeneous and complex, which certainly alters Aβ concentrations. Our simplified geometry also employs a blood sink term (−*βC*_*i*_) rather than solving the much more complex problem of (non-Newtonian) blood flow in a capillary. In addition, our model only considers one type of vessel: a capillary immediately surrounded by brain tissue. Larger vessels, including arteries and veins, are immediately surrounded by a perivascular space filled with flowing CSF [62]. Hence, Aβ clearance in the vicinity of these larger vessels likely involves interactions between the glymphatic system and BBB [42], rendering such simulations substantially more complex. We note that there is some evidence that capillaries may have a perivascular space [63], which we have neglected in this study. Lastly, our modeling of sleep and SD are highly idealized with several assumptions, including a constant rate of Aβ generation (*G*). Future studies should improve upon these numerous limitations and may even eventually perform mouse-specific simulations of geometry and vascular dynamics obtained *in vivo* using high-resolution optical imaging techniques such as two-photon microscopy.

Overall, this work introduces a mechanistic and biophysically grounded boundary condition for solute exchange via active transporters at the BBB, offering a valuable tool for future studies on drug delivery, brain waste clearance, and modeling pathogenesis of neurodegenerative diseases. Future simulations could probe active transport of molecules other than Aβ, including glucose, lactate, amino acids, or pharmaceuticals [4, 64]. Such simulations may generate predictions that guide future experiments. The considerable dependence on receptor number ratio and *β* suggest future experiments should seek to better quantify these important parameters. However, *β* arises as an idealization of blood flow, so future modeling studies could instead focus on extending our idealized axisymmetric model to more realistic two- or three-dimensional domains with explicit implementation of blood flow to improve understanding of BBB active transport.

## ACKNOWLEDGMENTS

Research reported in this publication was supported by the National Institute Of Biomedical Imaging And Bioengineering of the National Institutes of Health under Award Number R21EB036217. The content is solely the responsibility of the authors and does not necessarily represent the official views of the National Institutes of Health.

## DATA AVAILABILITY

The code used to generate the computational data and reproduce the figures of the manuscript will be deposited on Zenodo, an open research repository, upon acceptance of this manuscript.

## Supplemental Material

### I. NONDIMENSIONALIZATION

To non-dimensionalize the equations, we define the dimensionless variables as follows

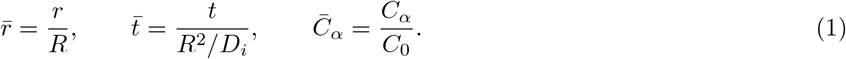

By substituting in these variables, Equations (18) to (19) of the main text become

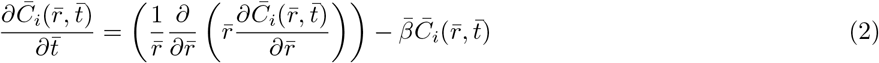

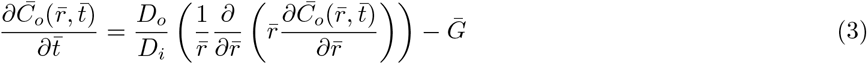

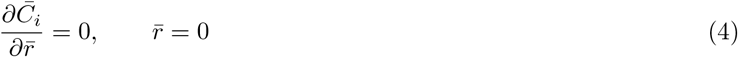

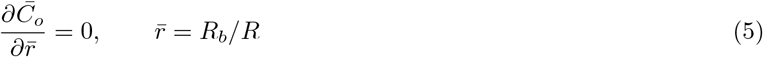

where

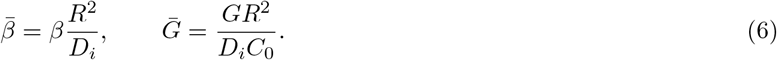

Additionally, the dimensionless form of the boundary condition introduced in Equation (27) of the main text becomes

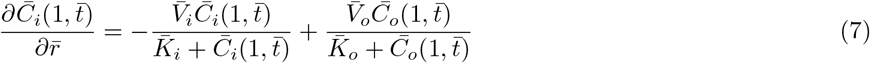

where

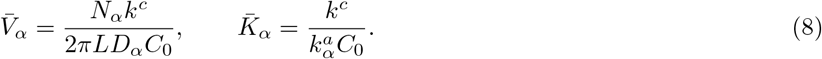

### II. LINEAR FORM OF THE BOUNDARY CONDITION

The boundary condition derived in this study (Equation (7)) accounts for various parameters of the BBB, including the number, size, and kinetics of the receptors. Although this adds complexity to both the derivation and the boundary condition itself, it results in more accurate and reliable outcomes. To examine the significance of these complexities, we performed the same simulations presented in Figure 4a of the main text using a simplified linear form of the boundary condition, as follows:

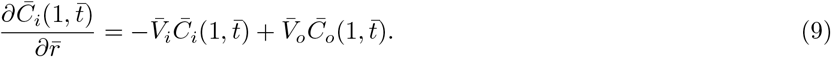

The results of this comparison, provided in Figure S1, reveal a significant difference between the results, highlighting the importance of the detailed boundary condition derived in this work. The linear boundary condition in Equation (9) neglects the saturating effects at the boundary due to high concentrations and the effect of the size of the receptors.

**FIG. S1.**
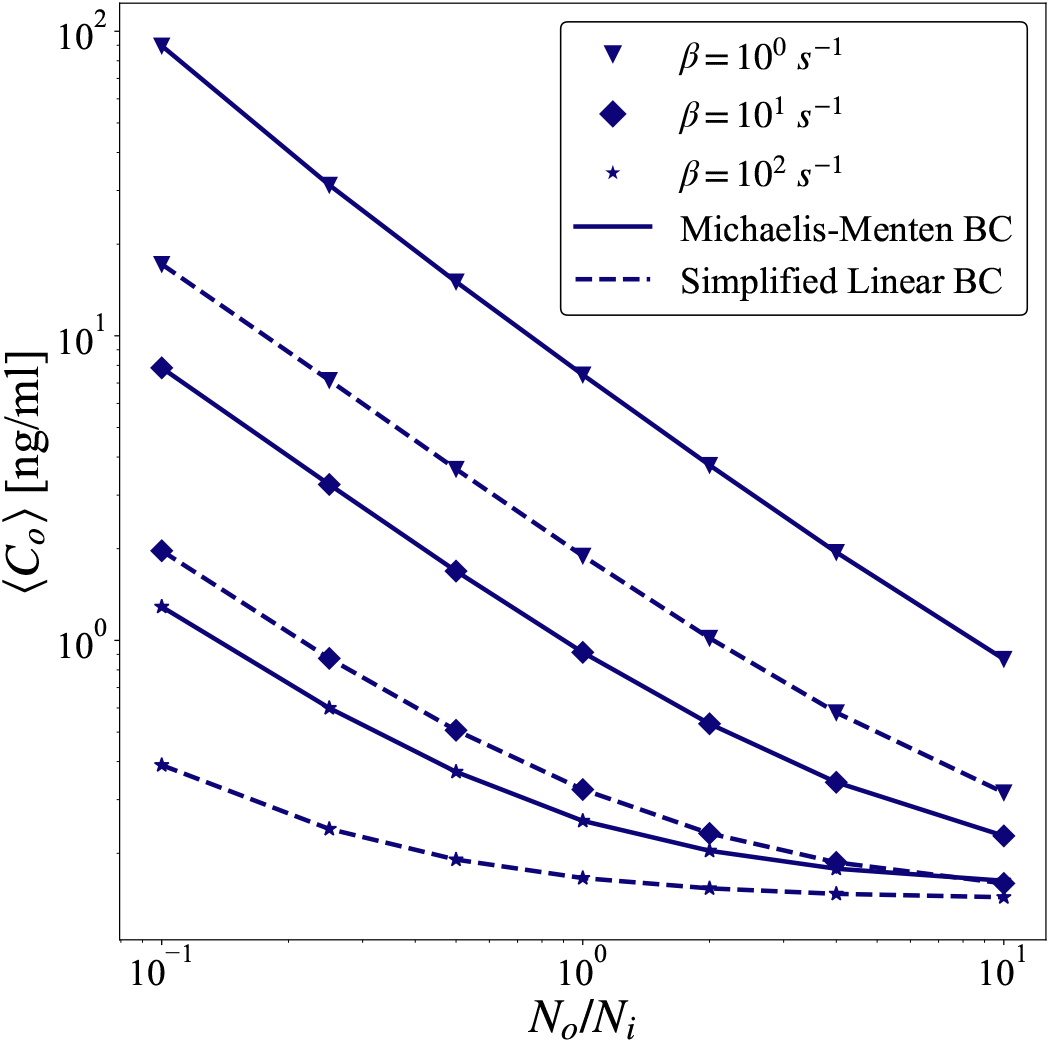
Averaged brain Aβ concentration as a function of blood sink coefficient β for the Michaelis-Menten boundary condition (derived in this work) and a simplified linear form of the boundary condition.

### III. NUMERICAL METHODS

We discretize space along the *r*-direction (indicated with a superscript) such that 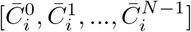 are on the interior of the vessel and 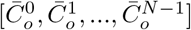 are in the brain tissue as shown in Figure S2. We define 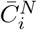 and 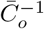 as ghost points outside of the specified regions to improve numerical implementation of boundary conditions. The discretized form of Equations (2-3) on the BBB boundary are as follows:

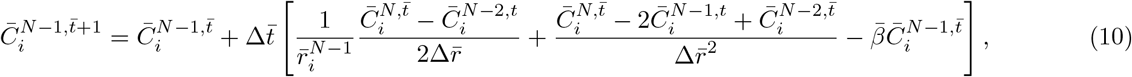

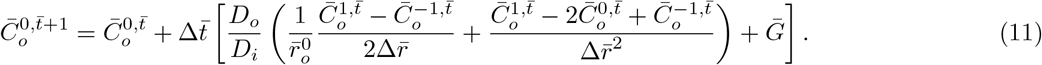

**FIG. S2.**
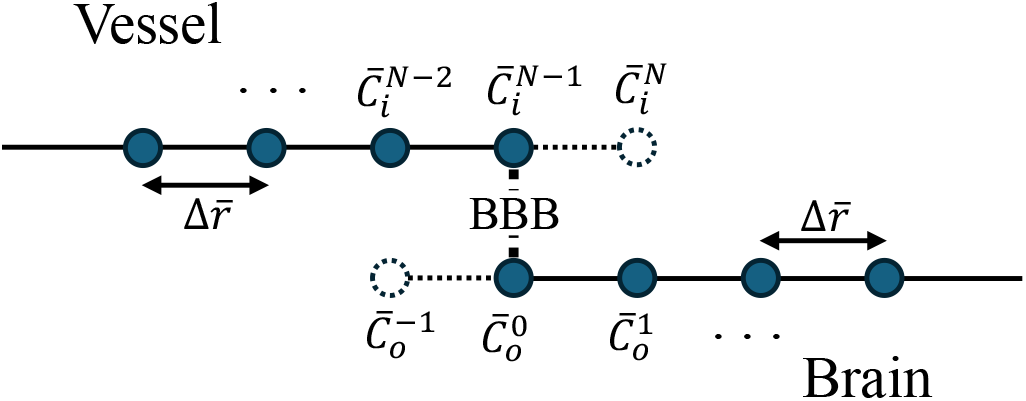
Discretization and BBB boundary-handling schematic.

We can use the boundary condition of Equations (18-19) to solve for 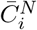 and 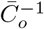. The discretized form of the BBB boundary condition is as follows:

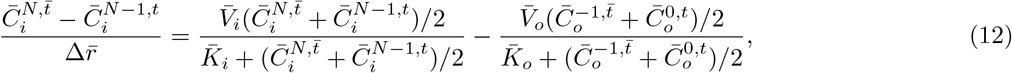

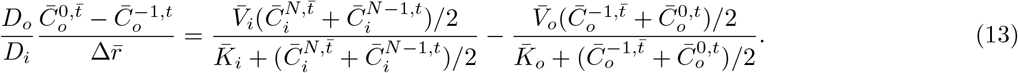

This is a non-linear system of two equations with two unknowns (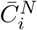 and 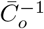) that we solve in every iteration in time to calculate the flux through the BBB boundary.

### IV. STEADY STATE ANALYTICAL SOLUTION

Recall our dimensionless system of partial differential equations

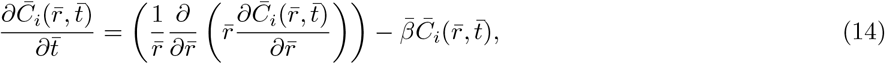

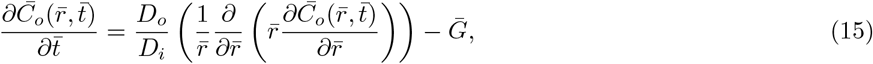

with boundary conditions

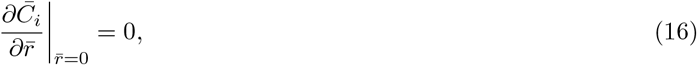

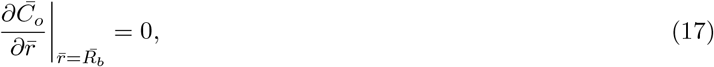

at the end points of the domain, and boundary conditions

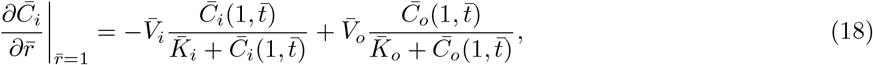

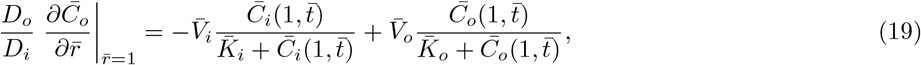

at the blood vessel interface in the interior. To solve for the steady state solution of this system of partial differential equations, we set the temporal derivative terms to zero and obtain the following system of ordinary differential equations,

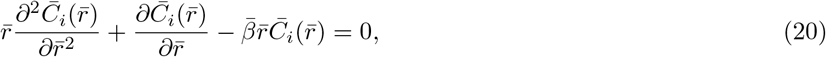

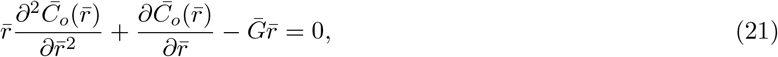

with the same boundary conditions as specified in Equations (16-19). Solving these equations with just the exterior boundary conditions in mind yields the following forms for the general solution of this system:

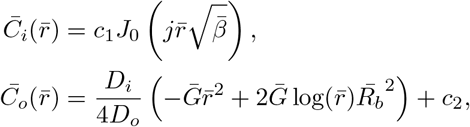

where *J*_0_(·) denotes the Bessel function of the first kind and *j*^2^ = −1. The remaining constants, *c*_1_ and *c*_2_ can be found by plugging these forms into boundary conditions (18-19), and solving accordingly.

We see that *c*_1_ can be readily solved for, as long as *V*_*o*_ *>* 0, by setting the right-hand sides of these equations to be equal, which yields

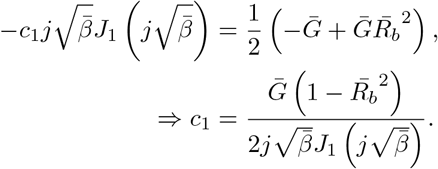

As a result, the solution in the blood vessel is, in terms of dimensional parameters,

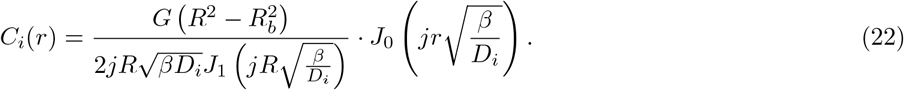

In this form, it is clear that this solution does not depend on the kinetic parameters of the transporter, particularly *N*_*i*_ and *N*_*o*_, as well *D*_*o*_.

